# SLIDE: Significant Latent Factor Interaction Discovery and Exploration across biological domains

**DOI:** 10.1101/2022.11.25.518001

**Authors:** Javad Rahimikollu, Hanxi Xiao, Anna E. Rosengart, Tracy Tabib, Paul Zdinak, Kun He, Xin Bing, Florentina Bunea, Marten Wegkamp, Amanda C. Poholek, Alok V Joglekar, Robert A Lafyatis, Jishnu Das

## Abstract

Modern multi-omic technologies can generate deep multi-scale profiles. However, differences in data modalities, multicollinearity of the data, and large numbers of irrelevant features make the analyses and integration of high-dimensional omic datasets challenging. Here, we present Significant Latent factor Interaction Discovery and Exploration (SLIDE), a first-in-class interpretable machine learning technique for identifying significant interacting latent factors underlying outcomes of interest from high-dimensional omic datasets. SLIDE makes no assumptions regarding data-generating mechanisms, comes with theoretical guarantees regarding identifiability of the latent factors/corresponding inference, outperforms/performs at least as well as state-of-the-art approaches in terms of prediction, and provides inference beyond prediction. Using SLIDE on scRNA-seq data from systemic sclerosis (SSc) patients, we first uncovered significant interacting latent factors underlying SSc pathogenesis. In addition to accurately predicting SSc severity and outperforming existing benchmarks, SLIDE uncovered significant factors that included well-elucidated altered transcriptomic states in myeloid cells and fibroblasts, an intriguing keratinocyte-centric signature validated by protein staining, and a novel mechanism involving altered HLA signaling in myeloid cells, that has support in genetic data. SLIDE also worked well on spatial transcriptomic data and was able to accurately identify significant interacting latent factors underlying immune cell partitioning by 3D location within lymph nodes. Finally, SLIDE leveraged paired scRNA-seq and TCR-seq data to elucidate latent factors underlying extents of clonal expansion of CD4 T cells in a nonobese diabetic model of T1D. The latent factors uncovered by SLIDE included well-known activation markers, inhibitory receptors and intracellular regulators of receptor signaling, but also honed in on several novel naïve and memory states that standard analyses missed. Overall, SLIDE is a versatile engine for biological discovery from modern multi-omic datasets.

## Introduction

Modern multi-omic technologies can generate deep multi-scale profiles. However, differences in data modalities, multicollinearity of the data, and large numbers of irrelevant features make the analyses and integration of high-dimensional omic datasets challenging. For example, multicollinearity can increase the variance of regression coefficients and lead to deflation of corresponding P values^1^. This is a significant barrier to meaningful inference in a regression setting for high-dimensional multi-collinear data. Further, human biological systems are complex, multi-factorial and organized hierarchically, with complex interaction rules at each hierarchy. A linear model is often inadequate at capturing relevant higher-order relationships in such a system. Finally, while recent methods developed by us^2-7^ and others^8-10^ have harnessed these high-dimensional multi-scale multi-modal datasets to predict different outcomes/groups of interest, they do not provide meaningful inference beyond prediction. In fact, approaches that do provide insights into the underlying mechanistic bases of outcome are tailored primarily for low-dimensional datasets, and often trade predictive power for inference^11^.

To address these, we present SLIDE, a novel data-distribution-free approach to analyze high-dimensional multi-omic datasets and uncover latent factors that drive the outcome of interest (Fig. 1a). SLIDE makes no assumptions regarding the distribution of the underlying data as it builds on a unique latent-factor regression framework developed by us^12, 13^. It takes into account an extremely large search space of relationships to converge on a very small subset of biologically relevant and actionable latent factors. Critically, SLIDE incorporates both linear and non-linear relationships, including complex hierarchical structures. It uncovers significant interacting latent factors in diverse contexts that span scales of organization from cellular/molecular phenotypes (e.g., extent of clonal expansion of CD4 T cells) to organismal phenotypes (e.g., disease severity of patients with diffuse systemic sclerosis). The discovery of these relationships is also coupled to rigorous FDR control, something that is extremely difficult to do in a large search space of non-linear/hierarchical relationships. Here, this is made possible using our unique analytical framework that creatively adapts ultra-modern methods for FDR control^14^. SLIDE comes with provable statistical guarantees regarding identifiability of the latent factors, corresponding inference of significant interacting latent factors. This is fundamentally different from several methods that have been published recently that rely on clever heuristics but do not have formal statistical guarantees or work only when strong biological priors are available. SLIDE has rigorous statistical guarantees, recapitulates known biological mechanisms and helps uncover novel biological mechanisms.

**Figure 1.**
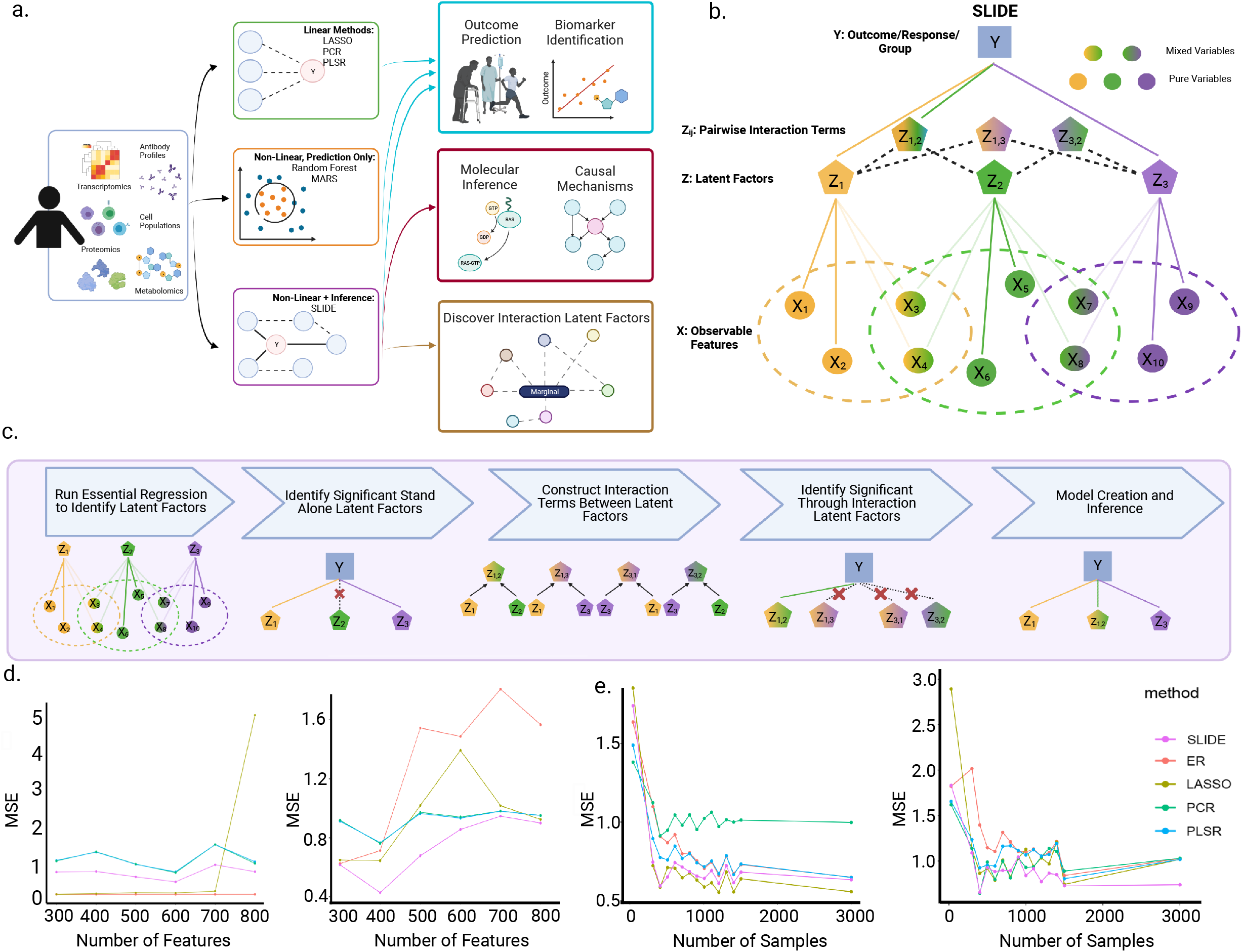
SLIDE – a novel interpretable machine learning method for Significant Latent Factor Interaction Discovery and exploration. a) Schematic illustrating the vast array of datasets on which SLIDE can be applied and the key advances over existing analytical frameworks for the analyses of these datasets b) Conceptual overview of the SLIDE algorithm. c) Schematic summarizing the implementation and different steps in SLIDE. d) Comparison of the predictive performance of ER, LASSO, PCR, PLSR and SLIDE on simulated datasets across a range of number of features without (left sub-panel) and with (right sub-panel) interaction terms. e) Comparison of the predictive performance of ER, LASSO, PCR, PLSR and SLIDE on simulated datasets across a range of sample sizes without (left sub-panel) and with (right sub-panel) interaction terms.

We tested the predictive performance of SLIDE on a range of datasets, and it outperformed several state-of-the-art approaches. Further, it provided novel inference not afforded by any existing approaches, thus being one of the only methods that simultaneously provides meaningful inference for high-dimensional data without compromising on predictive power. When analyzing datasets from SSc patients to elucidate the basis of SSc pathogenesis, SLIDE recovered altered transcriptomic states in myeloid cells and fibroblasts, a well elucidated basis of SSc disease severity^15-20^. But it also identified an unexplored keratinocyte-centric signature (validated by protein staining), and a novel mechanism involving an interaction between the altered transcriptomic states in myeloid cells and fibroblasts with HLA signaling in macrophages. This mechanism has strong support in recent genetic association analyses^21^. In the characterization of latent factors underlying clonal expansion of CD4 T cells, SLIDE recapitulated well known inhibitory receptors and markers of activation/exhaustion including Lag3, Pdcd1 (Pd1), Tigit and Cd200^22^ that standard differential expression (DE) analyses would have picked up. However, it also identified several novel markers that standard DE analyses would have missed. This includes novel gene sets and functional modules that have both significant linear and non-linear relationships to clonal expansion. Overall, SLIDE is a first-in-class approach for the discovery of significant interacting latent factors from high-dimensional multi-modal datasets that help infer novel biological mechanisms. It is an engine for biological discovery from modern multi-omic datasets.

### The SLIDE framework

SLIDE is an interpretable latent factor regression approach (Fig. 1b) that seeks to identify significant latent factors that capture linear and nonlinear relationships (up to pairwise interactions) between observed data (X, typically high-dimensional, multi-collinear) and the response/group/outcome of interest (Y) as presented in equations 1 and 2.

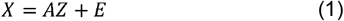

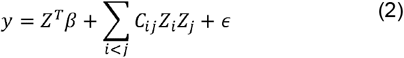

Given that *x*_*n*×*p*_ is a matrix of observables, where n and p represent the number of samples and features, respectively. The initial step (Fig. 1c) is to use our previously described LOVE approach^13^, to decompose the observable X into its constituent components of *A*_*P*×*K*_ and *Z*_*K*×*n*_, with an error term of *E*. Here, A is the allocation matrix and represents the membership of each feature to a latent factor. The *β* ∈ *R*^*K*×1^ and *C*_*ij*_ is the interaction effect between two latent variables of *Zi* and *Zj*. This approach comes with identifiability guarantees regarding inference of the latent factors without making any assumptions regarding data-generating mechanisms^13^.

The next step in SLIDE (Fig. 1c) is identifying significant latent factors using a multi-stage adaptation of an ultra-modern framework for FDR-controlled variable selection – knockoffs^14^. This approach is based on differences or lack thereof between true and fake (knockoff) variables. These knockoff variables should be orthogonal to the response variable. And these variables preserve the covariance structure (Σ). These variables are also approximately orthogonal to their original variables with deviation magnitude of 1-s. This means that the correlation between the original variable *Z*_*i*_ and the knockoff variable 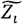 is 1-s, while the desired value is zero with s=1.

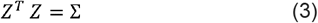

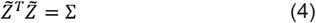

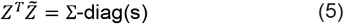

factors. Here, the variable *Z*_*i*_ is statistically significant if it considerably outperforms its knockoff While knockoffs have typically been used on observed variables, we used it on the latent 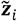 based on test statistics. In this study we used *W*_*j*_ as the test statistic of interest:

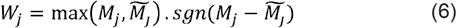

Here 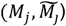 is defined as the maximum value of the L1 regularization hyperparameter *λ* corresponding to which the actual variable (*M*_*j*_) and its corresponding knockoff 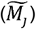 are hyperparameter *λ* indicates higher variable importance i.e., even with more stringent incorporated into the corresponding generalized linear model. A higher value of the regularization the feature stays on in the model. Thus, *W*_*j*_ (higher scores) will select for important latent factors with matched unimportant knockoffs.

Further, our adaptation of the knockoff approach is a multi-stage stage procedure (Fig. 1c). In stage 1, the latent factors are divided into smaller sets and the knockoff technique applied to each of these sets. This initial selection procedure helps shortlist putative significant latent factors for the entire dataset. In stage 2, these putative significant latent factors from each set undergo another round of selection via the Knockoff algorithm to converge on a set of standalone significant latent factors. We repeat these 2 stages multiple times to identify a set of stable standalone significant latent factors (Methods and Supplementary Methods).

In stage 3, we use hierarchical OR logic to identify significant interactors of the standalone significant latent factors i.e., each interaction term has at least one standalone significant latent factor. For each of the standalone significant latent factors, the interaction terms are identified by pairing it with the rest of the latent factors. The knockoff procedure is applied again to extract the significant interaction terms. In high dimensional datasets, where the number of features is much greater than the number of samples, the number of selected linear and non-linear latent factors might easily outnumber the number of observations. To account for this, we calculate the embedding of each significant latent variable and its subsequent interaction terms to a vector. Hierarchical information of the latent variable and its interactions is potentially preserved in these vectors. Therefore, the overall model quality can be attained by regressing the embedded features on the response variable with a smaller number of features than observables through cross-validation (Supplementary Methods).

First, using simulations (Methods), we compared the performance of SLIDE was compared to that of the state-of-the-art methods including ER^12^, LASSO^23^, Partial Least Squares Regression (PLSR)^24^, and Principal Components Regression (PCR)^25^ with and without interactions terms (Figs. 1d and 1e). SLIDE performs as well as state-of-the-art approaches when there are no interaction terms present (Figs. 1d and 1e). In the presence of interaction terms, it consistently outperforms these methods (Figs. 1d and 1e). Importantly, all approaches other than SLIDE and LASSO, use the full model (all features/clusters) for prediction. However, SLIDE only uses a small number of prioritized latent factors for prediction and yet achieves the same predictive power. It also provides inference not afforded by any other method including LASSO.

### SLIDE uncovers novel interacting latent factors that explain SSc pathogenesis

Using SLIDE, we first sought to discover interacting latent factors underlying SSc disease severity. We analyzed scRNA-seq data from 24 SSc subjects^15, 26^ across the severity spectrum (Fig. 2a), where disease severity was quantified using the Modified Rodnan Skin Score (MRSS). ScRNA-seq data from 24 SSc patients all using the same V2 chemistry was processed using a standard analytic pipeline. This consisted of alignment via *cellranger* and dimensionality reduction and clustering using Seurat (Methods). We identified 35 unique clusters (Fig. 2b). For downstream analyses, we filtered out clusters that contained less than 20 cells in any of the 24 subjects (Fig. 2b). For each of the retained clusters, we tried a range of feature engineering approaches and converged on a cell-type-specific pseudo-bulk average of the most variable genes, where variance was calculated in an unsupervised fashion without using the disease severity labels in any form (Methods). Next, we applied SLIDE on these cell-type-specific transcript abundances to predict SSc severity and infer corresponding significant interacting latent factors of outcome (Figs. S1a and S1b). SLIDE was able to accurately predict SSc severity and outperformed or 3/5 of our benchmarks – PLS^24^,, PCR^25^ and Phate-Regression (linear regression coupled to PHATE^27^) in terms of prediction accuracy (Figs. 2c and S1c). LASSO^23^ and ER^12^ (developed by us) were the only methods with comparable prediction performance (Fig. 2c). However, LASSO only identified a small set of predictive biomarkers that were uninformative of the actual molecular basis underlying SSc pathogenesis. On the other hand, SLIDE identified 9 significant latent factors that could be used to infer the mechanistic basis of SSc pathogenesis. Further, while the performance of SLIDE and ER were comparable, ER used the entire set of latent factors to predict outcome, while SLIDE only used 9. Thus, SLIDE provides the same predictive power as ER but stronger inference with fewer latent factors. Further, as a reference, if we used only the 9 most significant latent factors from ER (chosen based on the linear regression coefficients) for prediction, it would significantly underperform SLIDE (Fig S1d).

**Figure 2.**
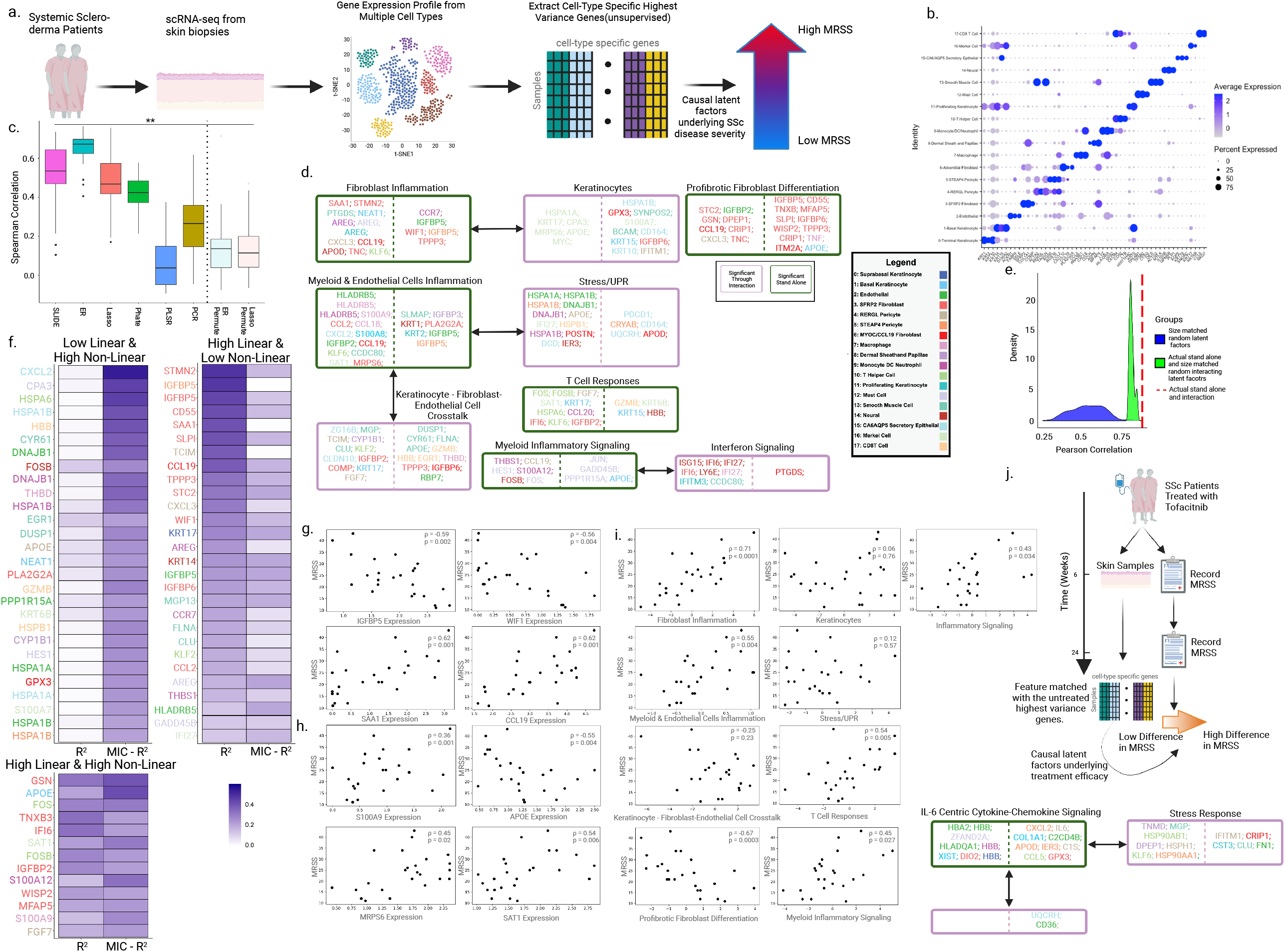
SLIDE uncovers novel interacting latent factors that explain SSc pathogenesis. a) Schematic summarizing scRNA-seq data from SSc patients used to infer mechanisms of SSc pathogenesis MRSS. b) Identities of the cellular clusters used in the analyses as defined by the top cell-type-specific DEGs. c) Spearman correlations between true MRSS and MRSS predicted using different methods – SLIDE (spec = 0.1), ER, LASSO, PLS, PCR and PHATE Regression. Model performance is measured in a k-fold cross-validation framework with permutation testing. ** indicates P < 0.01. d) Significant interacting latent factors identified by SLIDE. Green boxes denote significant standalone latent factors, and purple boxes denote significant interacting latent factors. The color of the gene corresponds to the cell type. Genes on the left and right of the dashed line have negative and positive correlations with MRSS respectively. e) Performance of the real model relative to a) the distribution of the performance of models built using size-matched random latent factors (blue) and b) the distribution of the performance of models built using the actual significant standalone latent factors and size-matched random interacting latent factors (green). f) Linear Spearman correlations and non-linear relationships (quantified using MIC) between key components of the latent factors and MRSS. g) Scatter plot of MRSS and gene expression of select key markers of SSc severity that have a significant linear relationship alone with MRSS. h) Scatter plot of MRSS and gene expression of select key markers of SSc severity that have a significant linear as well as a significant non-linear relationship with MRSS. i) Scatter plot illustrating the relationships between each significant latent factor (from d) and MRSS. j) Schematic summarizing a recent clinical trial where SSc patients are treated with tofacitinib. scRNA-seq data and changes in MRSS on treatment are available for these patients. Significant standalone and interacting latent factors underlying changes in MRSS on treatment are inferred.

The 9 latent factors uncovered by SLIDE spanned a range of cell-intrinsic and cell-extrinsic circuits (Fig. 2d). These factors encompassed altered transcriptomic states that have been characterized and are widely recognized to be critical in SSc pathogenesis. These states include modulated inflammatory states/signaling in myeloid cells and fibroblasts, including SFRP2 fibroblasts, well-known bases of SSc pathogenesis (Fig. 2d) ^15-20^. Other canonical mechanisms recapitulated by our model includes cross-talk between interferon signaling and myeloid inflammatory signaling (Fig. 2d)^15-20^. Key genes that contribute to these altered transcriptomic states include cytokines and chemokines (e.g., CCL19), signaling molecules (e.g., WIF1), interferon signaling genes (e.g., IGFBP5), components of mechano-transduction (e.g., THBS1) and alarmins/damage sensing molecules (e.g., S100A9). These agree well with previous studies by us and others^15-20^. However, in addition to recovering well-known mechanisms, we converged on several novel mechanisms. The first involves a previously unelucidated role of keratinocytes in SSc pathogenesis (Fig. 2d). We have recently validated this keratinocyte functional signature by protein staining (in a separate manuscript in revision). We also converged on another novel mechanism involving interactions (a key innovation of SLIDE) between altered myeloid/endothelial cell inflammation and keratinocyte-fibroblast-endothelial cell crosstalk. A key component of this interaction is the relationship between altered HLA signaling and modulated cytokine/chemokine/interferon signaling (Fig. 2d). While our study is the first to study this at the transcriptomic level, there is evidence for this mechanism in recent genetic studies^21^.

Interestingly, while we had 5 standalone significant latent factors, the 4 interacting latent factors provided additional information that was not encapsulated in the standalone factors. To formally test how relevant this information is, we first analyzed the predictive power of a size-matched set of random latent factors (Figs. 2e and S1e). The actual latent factors performed significantly better than the random size-matched set of latent factors (Figs. 2e and S1e). Next, we evaluated the quality of the interactors themselves by keeping the two actual standalone latent factors fixed but shuffling the interactors (i.e., choosing a size-matched set of random interactors for the actual standalone latent factors). Again, the performance of this model was significantly lower than the performance of the actual model (Figs. 2e and S1e). This is an especially stringent test as the standalone factors are highly informative themselves. Yet, not having the correct interacting latent factors would decrease the prediction and corresponding inference of mechanisms underlying SSc pathogenesis.

To further dissect these 9 significant latent factors (standalone and interacting), we tested the nature of relationships between the genes in these latent factors with MRSS. Interestingly, we observed that while some well-described/canonical markers of SSc severity such as CCL19, IGFBP5, WIF1, SAA1 and THBS1^15-20^, primarily in the SFPR2 fibroblasts and myeloid compartments, had significant high linear correlations (quantified using Spearman correlations) with MRSS, but almost no non-linear relationships with MRSS (Figs. 2f-2h). Further, several other genes such as APOE, S100A9 have both significant linear and non-linear relationships with MRSS (Figs. 2f-2h). Some of these are entirely novel, and others that have begun to be characterized in SSc^15-20^. Finally, several have only non-linear relationships with MRSS (Fig. 2f-2h). Most of these have been missed by previous approaches. Further, we evaluated the relationships of each of the 9 latent factors with MRSS (Fig. 2i) and found significant relationships between most individual latent factors and MRSS. This analysis suggests that SLIDE captures true context-specific biological group structure i.e., beyond the overall multivariate model, each individual context-specific group (latent factor) has meaningful information (both in terms of prediction and inference) regarding SSc pathogenesis (Fig. 2i). Canonical approaches focused on gene lists do not have group information at all, while pathway-centric approaches have group information that is context-independent and hence often irrelevant in specific scenarios.

To more rigorously test the actual causal nature of these identified latent factors, we moved to a “human perturbation experiment” made possible by the fact that 10 of these 24 subjects had recently been in a recent clinical trial with tofacitinib (tofa) (Fig. 2j)^28^. Tofa is a IL6/JAK inhibitor^29^ that led to a reduction in SSc severity of some patients but not others (Fig 2j). We used SLIDE to identify significant interacting latent factors that underlie the reduction in disease severity (change in MRSS from pre-to post). We hypothesized that if the identified latent factors were indeed causal, it would capture relevant tofa-centric mechanisms that led to a reduction in SSc severity (Fig. 2j). Further, we only had SSc scRNA-seq signatures 6 weeks post treatment i.e., it was an early snapshot. However, the measured change in disease severity was 24 weeks post treatment i.e., it reflected true response to outcome (Fig. 2j). SLIDE identified 3 significant interacting latent factors underlying the response to tofa. These included an IL6-centric cytokine-chemokine signaling latent factor in myeloid and fibroblasts and response to stress. Remarkably, despite not incorporating any biological priors, SLIDE very accurately honed in on the exact IL6-centric molecular mechanism^28^ (from thousands of transcriptomic features) underlying tofa treatment, demonstrating its power in identifying true causal mechanisms from complex high-dimensional datasets (Fig. 2j). And it was able to do so even at an early timepoint, demonstrating the sensitivity of SLIDE in capturing subtle changes in the course of treatment.

### SLIDE uncovers latent factors underlying immune cell partitioning by 3D localization

We then applied SLIDE to spatial transcriptomic datasets aiming to uncover latent factors underlying 3D spatial partitioning of immune cells. Spatial RNA-seq using 10X Visium was performed in a murine house dust mite model for studying allergy^30, 31^. Specifically, animals were treated intranasally with house dust mite for three consecutive days and mediastinal lymph nodes were isolated from these animals followed by spatial RNA-seq (Fig. 3a, Methods). We overlayed the clustering of the spatial regions with fluorescence microscopy images to designate spatial labels of border, central and intermediate zones (Fig. 3b). We also observed that the border regions were enriched for B-cells (blue) and the central regions for CD4 T cells (green) and dendritic cells (pink, DCs) (Fig. 3b). Since SLIDE does not make any assumptions regarding data-generating mechanisms, this tests the power of SLIDE on analyzing datasets generated using ultra-modern technologies.

**Figure 3.**
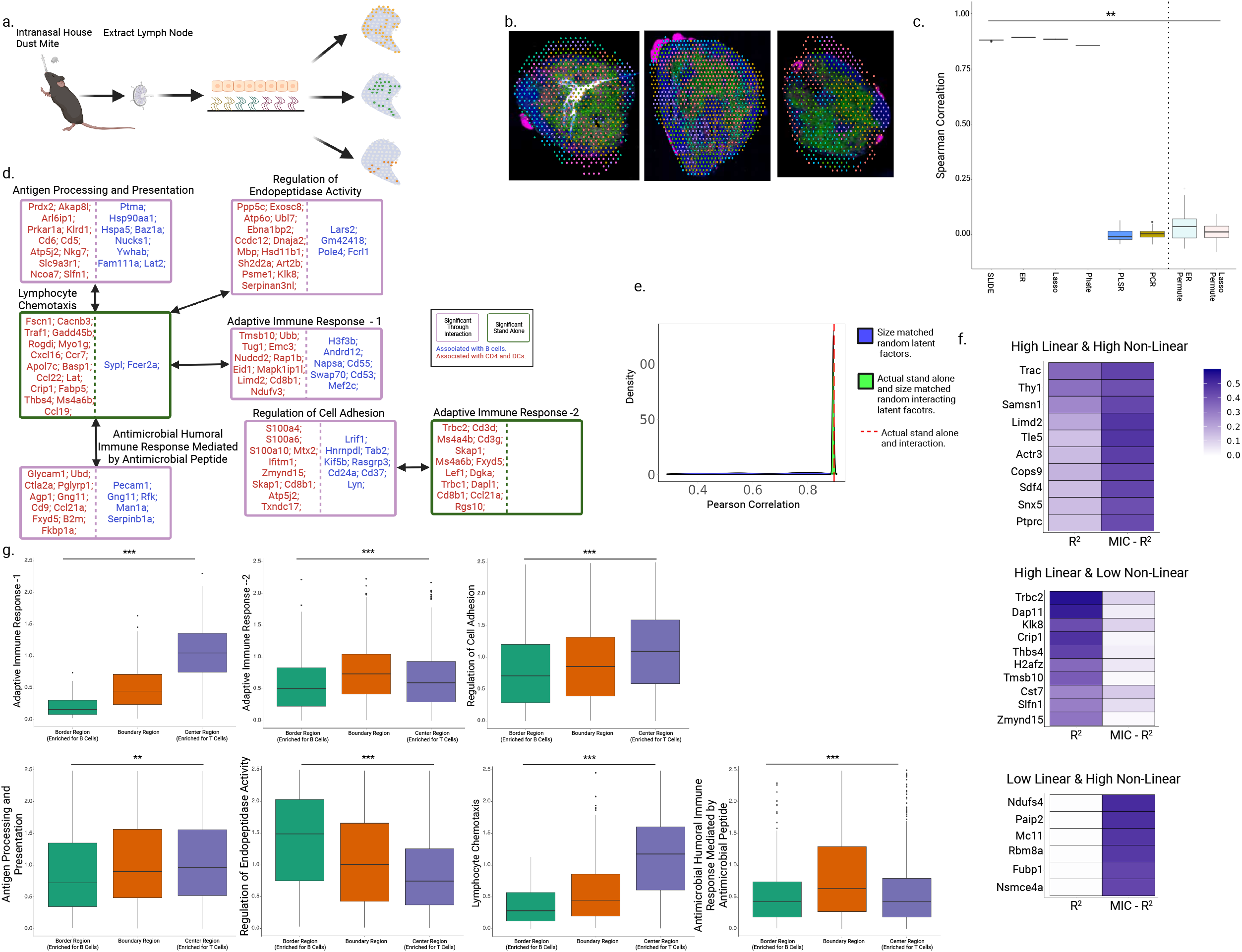
SLIDE uncovers latent factors underlying immune cell partitioning by 3D localization. a) Schematic summarizing scRNA-seq data from SSc patients used to infer mechanisms of SSc pathogenesis MRSS. b) KNN clustering of the spatial regions overlayed with microscopic images of three technical replicates of murine lymph node. Blue dye represents B-cells. Green dye represents CD4 T-cells and dendritic cells. Boundaries are categorized as boundary cells. c) Spearman correlations between true spatial region and spatial region predicted using different methods – SLIDE (spec = 0.1), ER, LASSO, PLS, PCR and PHATE Regression. Model performance is measured in a k-fold cross-validation framework with permutation testing. ** indicates P < 0.01. d) Significant interacting latent factors identified by SLIDE. Green boxes denote significant standalone latent factors, and purple boxes denote significant interacting latent factors. The color of the gene corresponds to the cell type. Genes on the left and right of the dashed line have negative and positive correlations with spatial region respectively. e) Performance of the real model relative to a) the distribution of the performance of models built using size-matched random latent factors (blue) and b) the distribution of the performance of models built using the actual significant standalone latent factors and size-matched random interacting latent factors (green). f) Linear Spearman correlations and non-linear relationships (quantified using MIC) between key components of the latent factors and spatial region. g) Boxplots illustrating the distributions of each latent factor across spatial regions.

SLIDE was able to accurately predict spatial labels, and outperformed PLS, PCR in terms of prediction accuracy (Figs. 3c and S2a-2c). LASSO, PhateRegression and ER had comparable prediction performance (Figs. 3c and S2c). However, LASSO and PhateRegression did not provide any insights into the nature of immune cell partitioning by spatial region. SLIDE identified 7 significant latent factors that could be used to infer the mechanistic basis of spatial partitioning of the immune cells. Further, while the performance of SLIDE and ER were comparable, ER used the entire set of latent factors to predict outcome, while SLIDE only used 7. Thus, SLIDE provides the same predictive power as ER but stronger inference with fewer latent factors. Further, as a reference, if we used only the 7 most significant latent factors, chosen by the highest linear regression coefficient, from ER for prediction, it would significantly underperform SLIDE (Fig. S2d).

Interestingly, although SLIDE was only given spatial region labels, the identified latent factors consisted of genes that mark B cells, CD4 T cells and DCs. This agrees well with the spatial partitioning of immune cells that the fluorescence microscopy images provide us (Fig. 3d), suggesting that SLIDE provides meaningful inference of the basis of immune cell partitioning by spatial regions in lymph nodes. The 7 latent factors uncovered by SLIDE included markers that represent multiple canonical functions including broad adaptive immune responses, antigen processing and presentation and specific humoral responses (Fig. 3d). As described earlier, we also analyzed the predictive power of a size-matched set of random latent factors (Figs. 3e and S2e). The actual latent factors performed significantly better than the random size-matched set of latent factors (Fig. 3e). We then evaluated the quality of the interactors themselves by keeping the two actual standalone latent factors fixed but shuffling the interactors (i.e., choosing a size-matched set of random interactors for the actual standalone latent factors). Again, the performance of this model was significantly lower than the performance of the actual model (Figs. 3e and S2e). Here too, not having the correct interacting latent factors would decrease the prediction and corresponding inference of mechanisms underlying clonal expansion.

While a number of genes in the significant latent factors of spatial partitioning had significant linear relationships with spatial label (Figs. 3f and S2f), several others had only non-linear relationships which would have been missed by traditional regression methods (Figs. 3f and S2f). The incorporation of these relationships strengthens corresponding inference. Next, we evaluated the relationships of each of the 7 latent factors with spatial region label (Fig. 3g) and found significant relationships between most individual latent factors and spatial region label (Fig. 3g). This provides additional support to our hypothesis that SLIDE captures true context-specific biological group structure i.e., beyond the overall multivariate model, each individual context-specific group (latent factor) has meaningful information (both in terms of prediction and inference) regarding the spatial region label of interest.

### SLIDE elucidates novel interacting latent factors underlying clonal expansion in T1D

Finally, we sought to analyze paired multi-omic datasets using SLIDE and uncover interacting latent factors underlying clonal expansion in T1D. Using paired scRNA-seq and TCR-seq data on islet-derived CD4 T cells in a non-obese diabetic (NOD) mouse model (Zdinak et al, accompanying manuscript), we labeled cells (Fig. 4a) based on the extent of their clonal expansion – single (1 clone), low (2-10 clones) or medium/high (>10 clones). This analysis has three additional facets in evaluating the capabilities of SLIDE beyond what the previous examples afford. First, it helps test the performance of SLIDE on paired multi-omic data (scRNA-seq and TCR-seq data) drawn from different distributions and with varying degree of sparsity. Second, we use a high-resolution cellular phenotype i.e., the clonality of individual CD4 T cells as the outcome labels of interest. This is different from the prior examples where we used organismal phenotypes (SSc severity) or 3D spatial location as the outcomes of interest. Third, it demonstrates the power of SLIDE in identifying interacting latent factors at a per-cell resolution in a disease model.

**Figure 4.**
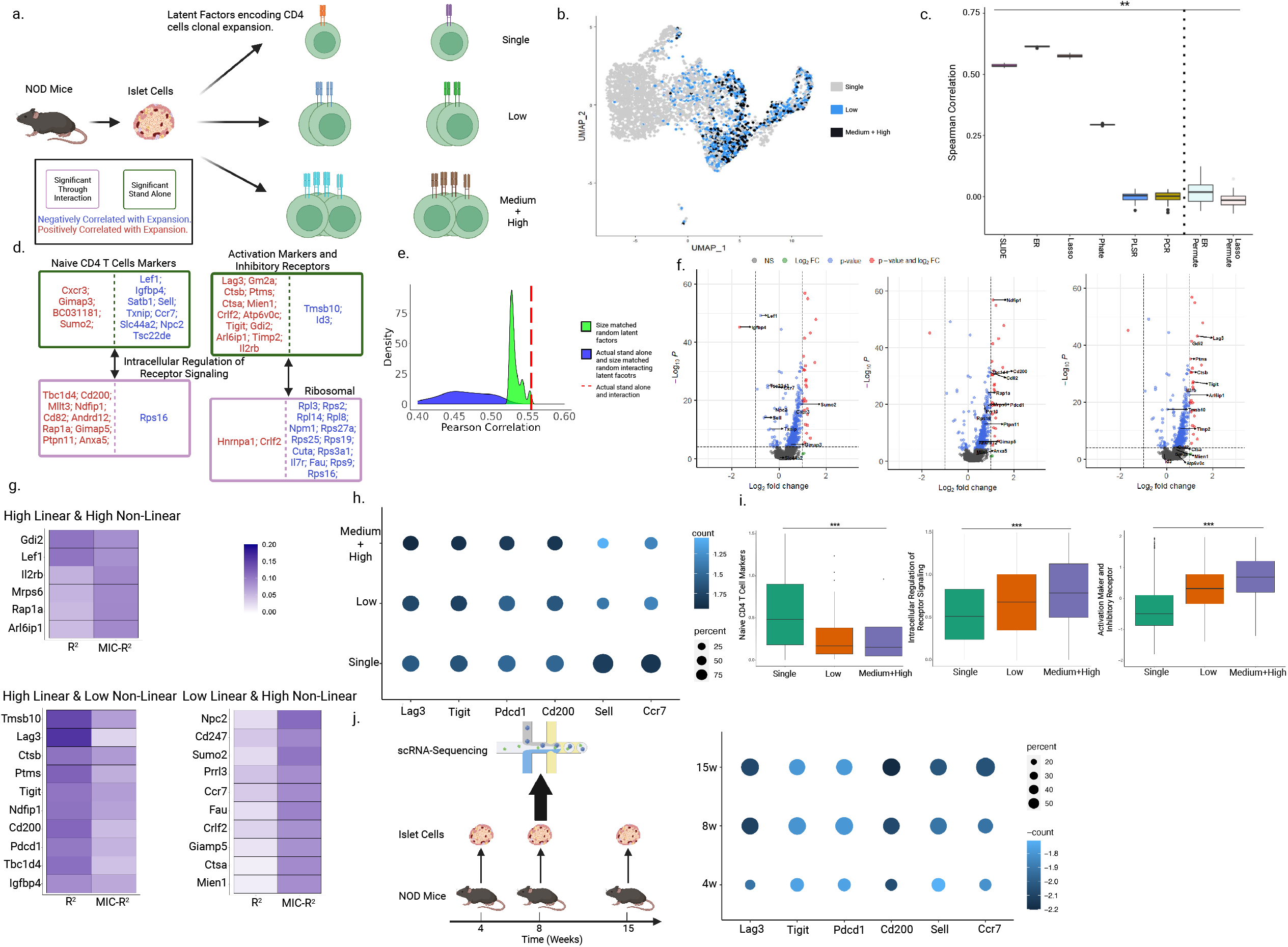
SLIDE elucidates novel interacting latent factors underlying clonal expansion in T1D. a) Schematic summarizing scRNA-seq and TCR-seq data from NOD mice used to infer mechanisms underlying clonal expansion of CD4 T cells. b) UMAP visualization of the three stages of clonal expansion. c) Spearman correlations between true stage of clonal expansion and stage of clonal expansion predicted using different methods – SLIDE (spec = 0.1), ER, LASSO, PLS, PCR and PHATE Regression. Model performance is measured in a k-fold cross-validation framework with permutation testing. ** indicates P < 0.01. d) Significant interacting latent factors identified by SLIDE. Green boxes denote significant standalone latent factors, and purple boxes denote significant interacting latent factors. The color of the gene corresponds to the cell type. Genes on the left and right of the dashed line have negative and positive correlations with extent of clonal expansion respectively. e) Performance of the real model relative to a) the distribution of the performance of models built using size-matched random latent factors (blue) and b) the distribution of the performance of models built using the actual significant standalone latent factors and size-matched random interacting latent factors (green). f) Volcano plots illustrating genes in the significant latent factors g) Linear Spearman correlations and non-linear relationships (quantified using MIC) between key components of the latent factors and extent of clonal expansion. h) Dot-plots illustrating frequency (circle size) and median expression (color intensity) of well-known markers of markers of T cell activation, exhaustion and inhibitory receptors at the 3 stages of clonal expansion. Frequency and expression calculated using data from our study. i) Boxplots illustrating the distributions of each latent factor across cells at the 3 different stages of clonal expansion. k) Dot-plots illustrating frequency (circle size) and median expression (color intensity) of well-known markers of markers of T cell activation, exhaustion and inhibitory receptors at the 3 stages of clonal expansion. Frequency and expression calculated using data from Unanue and colleagues. Schematic also summarizes design of study from Unanue and colleagues.

Interestingly, while there were some differences in transcriptomic profiles between cells at different stages of clonal expansion (Fig. 4b), there is significant heterogeneity in the profiles of individual cells at the same stage of clonal expansion (Fig. 4b). This necessitates the use of an approach like SLIDE that can converge on specific components of transcriptomic profiles that transcend intra-group heterogeneity and focus only on differences across cells at different stages of clonal expansion. SLIDE was able to accurately predict and infer extent of clonal expansion, and outperformed several benchmarks including PLS, PFR and Phate-Regression in terms of prediction accuracy (Figs. 4c and S3a-3c). LASSO and ER were the only methods with comparable prediction performance (Figs. 4c and S3c). However, LASSO only identified a small set of predictive biomarkers that were uninformative of the actual molecular basis for difference in clonal expansion across the different cells. On the other hand, SLIDE identified 4 significant latent factors that could be used to infer the mechanistic basis of differences in clonal expansion. Further, while the performance of SLIDE and ER were comparable, ER used the entire set of latent factors to predict outcome, while SLIDE only used 4. Thus, SLIDE provides the same predictive power as ER but stronger inference with fewer latent factors. It also identified non-linear relationships underlying clonal expansion (Fig. 4d). Further, as a reference, if we used only the 4 most significant latent factors, chosen by the highest linear regression coefficient, from ER for prediction, it would significantly underperform SLIDE (Fig S3d). The 4 latent factors uncovered by SLIDE included a) markers of Naive CD4 T cells, b) activation markers and inhibitory receptors, c) a latent factor that captured intracellular regulation of receptor signaling and d) ribosomal proteins (Fig. 4d). As earlier, we tested if the interactions of (a) with (c) and (b) with (d) provided better prediction and additional inference that (a) and (b) alone wouldn’t provide. Consistent with what we had observed in the earlier analyses, the actual latent factors performed significantly better than the random size-matched set of latent factors (Figs. 4e and S3e). Next, we evaluated the quality of the interactors themselves by keeping the two actual standalone latent factors fixed but shuffling the interactors (i.e., choosing a size-matched set of random interactors for the actual standalone latent factors). Again, the performance of this model was significantly lower than the performance of the actual model (Figs. 4e and S3e). In this case too, not having the correct interacting latent factors would decrease the prediction and corresponding inference of mechanisms underlying clonal expansion.

Importantly, the 4 significant latent factors included well-known inhibitory receptors and markers of clonal expansion/exhaustion including Lag3, Pdcd1 (Pd1), and Tigit^22^ that standard differential expression (DE) analyses would have picked up (Fig. 4f). As expected, SLIDE grouped these inhibitory receptors in one latent factor, suggesting potential convergent mechanisms. Interestingly, the intracellular signaling regulation latent factor (c) also contained Ndfip1^32^, which was shown to induce apoptosis in self-reactive T cells. The association of other potential mediators of apoptosis such as Anxa5^33^, suggests a different pathway of action than the inhibitory receptor latent factor. A traditional DE analysis simply returned a list of DEGs, but SLIDE’s grouping of genes with convergent functions led to the identification of inhibitory receptors and intracellular restriction on proliferation as two parallel mechanisms at work in clonally expanded T cells.

However, it also identified several novel markers that standard DE analyses would have missed. This includes genes that have both significant linear and non-linear relationships (at the univariate level) to clonal expansion. While the standard DE analyses would have picked up only the linear relationships, SLIDE can identify both (Fig. 4g). Of particular interest, is Ccr7, which is elevated in naïve T cells, as well as memory T cells. Co-expression of Ccr7 with Sell and Lef1 are hallmarks of naïve T cells^34^, confirming that unexpanded CD4+ T cells are naïve in their phenotype. Overall, the significant interacting latent factors are far better at capturing the molecular basis of clonal expansion than individual canonical markers previously reported in the literature (Figs. 4h and i). While the individual markers all recapitulate well-known trends (Lag3, Tigit, Pdcd1 and Cd200 are higher in expanded cells while Sell and Ccr7 are lower in unexpanded cells), the individual univariate relationships are weak (Fig. 4h). However, the significant latent factors correctly encapsulate additive effects and provide clear stratification by extent of clonal expansion. This analysis too provides support to our hypothesis that SLIDE captures true context-specific biological group structure i.e., beyond the overall multivariate model, each individual context-specific group (latent factor) has meaningful information (both in terms of prediction and inference) regarding the extent of clonal expansion.

Finally, to further validate and contextualize discoveries made by SLIDE, we analyzed scRNA-seq data from an independent recent study^35^ that had identified cell-specific markers of disease progression in T1D. Several key markers indeed followed the same trend (Fig. 4j) in the other study, confirming the robustness of our findings. However, importantly, not all relationships were the same as our markers reflect the extent of clonal expansion (a cellular phenotype) while the study identified markers of T1D disease progression (a related but different organismal phenotype). Overall, SLIDE is able to identify highly context-specific markers of clonal expansion of CD4 T cells in a NOD model of T1D.

## Discussion

With the rapid growth of technologies for deep profiling, there has been a deluge of high-dimensional datasets quantifying different facets of multi-scale multi-modal biological responses. There has been a significant investment in the development and application of statistical techniques for these high-dimensional datasets, but most of the methods have focused solely on prediction. Methods such as black-box deep learning approaches or classification/regression techniques based on higher-order embeddings are often highly accurate but are typically uninterpretable. Thus, they are useful in contexts such as clinical decision making (e.g., predicting disease severity/outcome), but provide very little or no insights into actual mechanisms of complex molecular, cellular or organismal phenotypes.

To address these key challenges, here we present SLIDE, an interpretable latent factor regression-based machine learning approach for ubiquitous biological discovery from high-dimensional multi-omic datasets. Unlike recent approaches that employ clever heuristics but lack statistical guarantees, SLIDE comes with rigorous guarantees regarding identifiability of the latent factors and corresponding inference. SLIDE can take into account complex non-linear and hierarchical relationships. The identifiability guarantees of SLIDE given it a significant edge over other modern techniques (e.g., variational autoencoders) that incorporate non-linear relationships, but are very sensitive to parameter initialization^1^. Further, most current state-of-the-art methods attempt to control FDR primarily through cross-validation and permutation testing. In addition to employing cross-validation and permutation testing, SLIDE creatively adapts knockoffs, an ultra-modern method for FDR control to identify significant standalone and interacting latent variables and their corresponding interactors. Also, to handle the complexity of high-dimensional datasets, SLIDE uses a multi-stage adaptation of knockoffs that includes runtime optimization feature space partitioning. SLIDE is also compatible with all pre-processing and batch-effect correction methods and/or technological-platform-specific analysis tools, because it makes no assumptions regarding data-generating mechanisms. Finally, SLIDE provides prediction better than or comparable to several state-of-the-art approaches, and inference that none of these techniques provide. Thus, unlike most methods, SLIDE provides inference in addition to, and not at the cost of predictive performance.

In addition to the wide array of conceptual and methodological innovation, SLIE also uncovers several novel biological mechanisms. In the analyses of transcriptomic datasets from SSc patients, in addition to recapitulating previously published markers^15-20^, we discovered several novel functional states underlying SSc pathogenesis. These include a previously unappreciated role of keratinocytes and a novel HLA-centric latent factor and its cross talk to myeloid and fibroblast cells. In the analyses of latent factors underlying clonal expansion of CD4 T cells in T1D, SLIDE recapitulated well-known activation markers and inhibitory receptors^22^, but converged on several novel signatures of naïve and memory states. Critically, the inference is multi-scale – both at the level of the overall model and individual context-specific groups (latent factors). Canonical approaches focused on gene lists do not have group information at all, while pathway-centric approaches have group information that is context-independent and hence often irrelevant in specific scenarios. This strengthens the interpretability of the model and provides more accurate guidance for potential downstream analyses and experimentation. Thus, SLIDE is truly a first-in-class interpretable machine learning framework for biological discovery.

## Methods

### Initial Latent Factor Discovery (LFD) Framework

The same LFD framework is employed for all three analyses presented in this work considering all linear relationship between latent factors and the corresponding dependent variable. The selection of significant standalone and interacting latent factors for each dataset using SLIDE is described in detail below. We first perform FDR thresholding on the covariance matrix, followed by the optimization of two key hyperparameters (delta and lambda) using *k* fold cross validation. The delta parameter controls the number of latent factors, while the lambda parameter has an impact on the sparsity of allocation matrix of latent factors. Considering the large and continuous search space of delta and lambda, we perform the search in multiple steps and ranges. The first step of the framework performs a coarse grid search of delta in four different numerical ranges: 0 – 0.001, 0.001 – 0.01, 0.01 – 0.1 and 0.1 – 1. The subsequent models utilize the most optimal delta within each range and identify the final delta using cross-validation and permutation testing. Once an optimal delta has been identified, lambda is tuned using a coarse grid search coupled to cross-validation and permutation testing.

### Simulations

The simulation aimed to evaluate model performance on different values of feature size (p) and sample size (n) with and without interaction terms. The numbers of latent variables (K), significant standalone and significant interacting latent variables were set 100, 2 and 5 respectively. For the models with no interactions, the number of interacting latent factors was set to 0. For varying features, we fixed the number of observations at 300. For simulations that have a varying n, we fixed the number of features to 1,000. To generate the simulated data, R package mvnorm was utilized to randomly generate datapoints K times, where K represents the number of latent factors.

For the model with interaction terms, the dependent variable is generated using equation 2 with coefficients generated randomly from standard normal distributions.

### Analyses of transcriptomic profiles in SSc patients

We analyzed scRNA-seq data from 24 previously described SSc patients where 10 of these patients were treated with Tofacitnib for 24 weeks. Standard 10X Genomics sequencing pipeline was used including *cellranger*. The aligned samples were then normalized and clustered with Seurat. We used R (version 4.00) for the data analysis. The Seurat (R package version 3.2.2) was adopted for cell population identification and visualization. To transform the 24 scRNA-seq samples into the pseudobulk fashion, we utilized the “Average Expression Function” to reduce the cell dimension by calculating the average expression of each gene across all cells in each cell-type specific cluster per patient. Clusters with fewer than 20 cells on average across all patients were excluded. 18 clusters passed the filtration process and the top 50 highest variance genes across patients of each cluster were chosen in an unsupervised manner as features for the subsequent analysis.

Post pre-processing, for the 24 untreated samples, the input data for the analysis is a sample by gene matrix with dimensions of 24 by 804. Using the LFD framework with 10-fold CV and 20 replicates, optimal delta and lambda values were identified (Figs S1a and S1b). With optimal parameter values set at 0.6 and 1 for delta and lambda respectively, the final model produced 120 latent factors. We then used SLIDE to identify the significant standalone latent factors using the iterative knockoff procedure as described above. Corresponding parameters of SPEC (a frequency-based parameter to quantify the stability of stages 1-2 of the multi-stage knockoff approach), FDR, F (feature split size) are set to 0.3, 0.1 and 100, respectively. The analysis resulted in 5 significant standalone factors and 4 significant interacting latent factors. 5-fold cross-validation with 50 replicates was used to compare the predictive power SLIDE with ER, LASSO, PHATE, PLSR, and PCR. Since PHATE is an unsupervised approach, we coupled it to a standard regression model (i.e., we ran Regression on PHATE1 and PHATE2). Glmnet and PhateR packages were used to build the LASSO and PHATE models, respectively. The PLS package was employed for PCR and PLSR model construction.

To assess linear and non-linear relationships of genes in the significant latent factors with MRSS, we used Spearman correlation and maximal information coefficient (MIC) respectively.

For the tofa-treated SSc analysis, post quality control, normalization and clustering using Seurat, we matched the cluster identities to the untreated samples. To transform the scRNA-seq dataset to the pseudo-bulk fashion, we used the same averaging calculation mentioned above and feature matched with the untreated analysis resulted in the input sample by gene matrix with dimensions of 10 and 728. Using the LFD framework, and LOOCV, a delta of 0.009 (175 latent factors) and a Lamba of 1 were identified as optimal hyperparameters. SLIDE was then applied to identify significant standalone and interacting latent factors. The SLIDE parameters of SPEC, FDR, and F were set to 0.3, 0.1 and 100 respectively. The final SLIDE model produced 1 significant standalone latent factor and 3 significant interacting latent factors.

### Analyses of spatial RNA-seq data in lymph nodes

25 μg of LPS-low HDM (Stallergenes-Greer Pharmaceuticals) in 25 μl of sterile 1X PBS was delivered intranasally under anesthesia to C57BL/6 mice (Jax Laboratory) daily for 3 days. Mediastinal lymph nodes (mLN) were isolated on day 4, snap frozen and embedded in chilled optimal cutting temperature (OCT) compound (Tissue-Tek) on dry ice, and stored at -80 °C. MLN samples were cryosectioned (10 μm) at -20 °C on a cryostat (Leica) and mounted directly onto the 6.5 × 6.5 mm capture areas of a single Visium Spatial Gene Expression slide (10X Genomics). The slides were sealed in individual 50 ml Falcon tube at -80 °C until further processing according to the manufacturer’s protocol (10x Genomics). The Visium Spatial Tissue Optimization Slide & Reagent kit (10x Genomics) was used to determine optimal permeabilization timing of 18 min. Immunofluorescence staining was done using “Methanol Fixation, Immunofluorescence Staining & Imaging for Visium Spatial Protocols (CG000312)”. Slides were stained with anti-B220 eFluor 450 (RA3-6B2, eBioscience), anti-CD4 Alexa Fluor 488 (G.K.1.5, Biolegend) and anti-CD11c Biotin (N418, Biolegend) followed by secondary detection with streptavidin Alexa Fluor-647. Images were acquired using EVOS™ M7000 Imaging System (AMF7000) under the Visium assay mode. Tissue sections were then permeabilized, and mRNA molecules within cells captured by poly (dT) sequence on the slide surface, followed by on slide reverse transcription to generate cDNA. cDNA is amplified and further processed into sequencing libraries according to the manufacturer’s protocol (10X Genomics). Libraries were sequenced on an Illumina Nextseq2000 at 50,000 read pairs per spot covered by tissue. Sequencing results were initially processed by *spaceranger* (10X Genomics) to align sequencing data with the image. Sequencing results were initially processed by *spaceranger* (10X Genomics) to align sequencing data with the image. Here, we filtered the genes more than 900 zeros threshold which resulted in matrix of 1932 genes by 3779 regions.

Seurat was employed for the quality control, normalization and clustering analyses. By overlaying the UMAP clusters and the fluorescent microscopy plot (Fig 3b), the UMAP clusters were assigned to the three different cell types: B-Cells, Boundary Cells and CD4/DCs. The regions that belong to the same cell types were pooled across the technical replicates. Genes that are not expressed in at least 900 regions were filtered to control the sparsity. Using the LFD framework with 10-fold cross-validation and 20 replications the optimal value for delta parameter was obtained as 0.049 with 21 latent variables (Fig. S2a). As described previously, 10-fold cross-validation with 20 replications was performed for optimal parameter tuning (Figs. S2a and S2b). SLIDE was applied to identify factors underline immune cell partitioning by spatial localization. We set an FDR threshold at 0.1 for each knockoff replicate and F = 21 for this dataset. 2 significant standalone latent factors and 5 significant interacting latent factors were identified.

### scRNA-seq and TCR-seq of islet infiltrating CD4 T cells

NOD mice were euthanized by CO2 asphyxiation and immediately dissected for pancreas perfusion. Pancreas perfusion was performed under a dissecting Zeiss microscope. Pancreatic duct was clamped using surgical clamps and 3 ml of 600 U/ml Collagenase dissolved in HBSS was injected using a 30G needle. Perfused pancreata were harvested and incubated at 37°C for 30 min. After the incubation, HBSS with R10 was added to quench collagenase. After washing twice with HBSS+R10, the tissue was plated on a 10 cm plate, individual islets were picked using a micropipettor. Islets were then incubated in dissociation buffer, centrifuged and, resuspended in the staining mix (1:500 dilution of anti-Thy1.2-BV605 + 1:500 dilution of Live/Dead-APC-Cy7, and 1:100 dilution of cell hashing anti-mCD45 TotalSeq-C antibodies (Biolegend)). After staining, the cells were resuspended in PBS+0.04% BSA and sorted on BD FACS Aria III sorter. After sorting the cells, they were counted and processed for scRNAseq. Cells were processing using 10x 5’ single cell gene expression kit v3 in a Chromium controller according to the manufacturer’s protocols. V(D)J enrichment was done using the single cell 5’ VDJ enrichment kit according to the manufacturer’s protocols. Libraries were sequenced on HiSeq4000 (Novogene Inc) with a 70:20:10 mix for gene expression:VDJ:hashing libraries as explained in greater detail in accompanying manuscript (Zdinak et al).

Sequence data were downloaded and aligned to the mouse genome (Mm10) using *cellranger* (10x Genomics Inc). TCR annotation was performed using *cellranger vdj* using mouse GRCm38 assembly. All three timepoints were sequenced and processed separately. Cellranger and cellranger vdj output files were used as inputs in Seurat for normalization, scaling, and dimensionality reduction. The packaged scRepertoire was used for TCR clonotype calling and analyses. The data were normalized using NormalizeData and scaled using ScaleData functions in Seurat. The scRepertoire functions combineTCR and combineExpression were used to add TCR clonotypes to each cell. HTODemux function in Seurat was used to demultiplex cell hashes and assign the correct mouse identity to each cell. At this point, all three timepoints were merged in Seurat using the merge function. After merging, integration was done using FindIntegrationAnchors and IntegrateData functions. Principle component analysis was performed using RunPCA. Top 20 principal components were used for Uniform Manifold Approximation and Projection, followed by cluster identification using FindNeighbors and FindClusters. CD4+ T cells were subsetted using FeatureScatter and CellSelector functions, and reclustered. Cluster markers were defined by FindAllMarkers function. Clonotype data were sorted according to expansion and exported as a csv file. UMAP representations with clonotypes were generated using highlightClonotypes function in scRepertoire. Differentially expressed genes were identified using FindMarkers function using DESeq2 statistics and represented using EnhancedVolcano function. After obtaining scRNA-seq and TCR-seq data on islet-derived cells in a non-obese diabetic (NOD) mouse model and analyzed data through the standard 10X Genomics pipeline. We labeled cells based on their clonal expansion stages followed by the post processing of the scRNA-seq data in R (4.1.0) using Seurat (R package, version 3.2.2). The columns representing genes and rows representing cells are filtered based on 1200 threshold, meaning that if the sparsity exceeds 1200, the cell row or gene column will be removed. The SLIDE input matrix was finalized with 1776 genes and 2482 cells.

The LFD framework is first utilized to discover latent factors. The input data is a cell by gene matrix, consisting of 2,484 cells and 1,776 genes. As described previously, 10-fold cross-validation with 20 replications was performed for optimal parameter tuning (Figs. S2a and b). The final model constructed by the LFD framework using delta as 0.0912 and lambda as 1 discovered 40 latent factors. We then performed SLIDE to identify significant interacting latent factors underlying differences in clonal expansion in CD4 T cells. We set the SLIDE parameter SPEC at 0.2, FDR at 0.1 and feature partition size at 40, resulting in the identification of two significant standalone and two significant interacting latent factors.

## Code and Data Availability

All code and data supporting all analyses in the manuscript are available at https://github.com/jishnu-lab/SLIDEpre and at https://github.com/jishnu-lab/SLIDE.

## Supplementary Figure Legends

**Figure S1. Accompanies Figure 2**

a) Spearman correlations between true MRSS and the predicted MRSS using different delta values in the LFD framework.

b) Spearman correlations between true MRSS and the predicted MRSS using different lambda values in the LFD framework.

c) Spearman correlations between true MRSS and MRSS predicted using different methods – SLIDE (spec = 0.3), ER, LASSO, PLS, PCR and PHATE Regression. Model performance is measured in a k-fold cross-validation framework with permutation testing. ** indicates P < 0.01.

d) Spearman correlations between true MRSS and MRSS predicted by ER (size-matched latent factors) and SLIDE.

e) Performance of the real model relative to a) the distribution of the performance of models built using size-matched random latent factors (blue) and b) the distribution of the performance of models built using the actual significant standalone latent factors and size-matched random interacting latent factors (green).

**Figure S2. Accompanies Figure 3**

a) Spearman correlations between true cell type region partitioning and predicted cell type region partitioning using different delta values in the LFD framework.

b) Spearman correlations between true cell type region partitioning and predicted cell type region partitioning different lambda values in the LFD framework.

c) Spearman correlation between true cell type region partitioning and cell type region partitioning predicted using different methods – SLIDE (spec = 0.2), ER, LASSO, PLS, PCR and PHATE Regression. Model performance is measured in a k-fold cross-validation framework with permutation testing. ** indicates P < 0.01

d) Spearman correlations between true MRSS and MRSS predicted by ER (size-matched latent factors) and SLIDE.

e) Performance of the real model relative to a) the distribution of the performance of models built using size-matched random latent factors (blue) and b) the distribution of the performance of models built using the actual significant standalone latent factors and size-matched random interacting latent factors (green).

f) Linear Spearman correlations and non-linear relationships (quantified using MIC) between key components of the latent factors and spatial region.

**Figure S3. Accompanies Figure 4**

a) Spearman correlation between true CD4 clonal expansion and predicted CD4 clonal expansion using different delta values in the LFD framework.

b) Spearman correlation between true CD4 clonal expansion and predicted CD4 clonal expansion different lambda values in the LFD framework.

c) Spearman correlation between true clonal expansion and clonal expansion predicted using different methods – SLIDE (spec = 0.3), ER, LASSO, PLS, PCR. Model performance is measured in a k-fold cross-validation framework with permutation testing. ** indicates P < 0.01

d) Spearman correlations between true MRSS and MRSS predicted by ER (size-matched latent factors) and SLIDE.

e) Performance of the real model relative to a) the distribution of the performance of models built using size-matched random latent factors (blue) and b) the distribution of the performance of models built using the actual significant standalone latent factors and size-matched random interacting latent factors (green).

f) Linear Spearman correlations and non-linear relationships (quantified using MIC) between key components of the latent factors and CD4 clonal expansion.

**Supplementary Methods – Algorithmic overview of SLIDE**

