## Supplementary figures and images for "SLIDE: Significant Latent Factor Interaction Discovery and Exploration across biological domains"

### Supplementary Figures

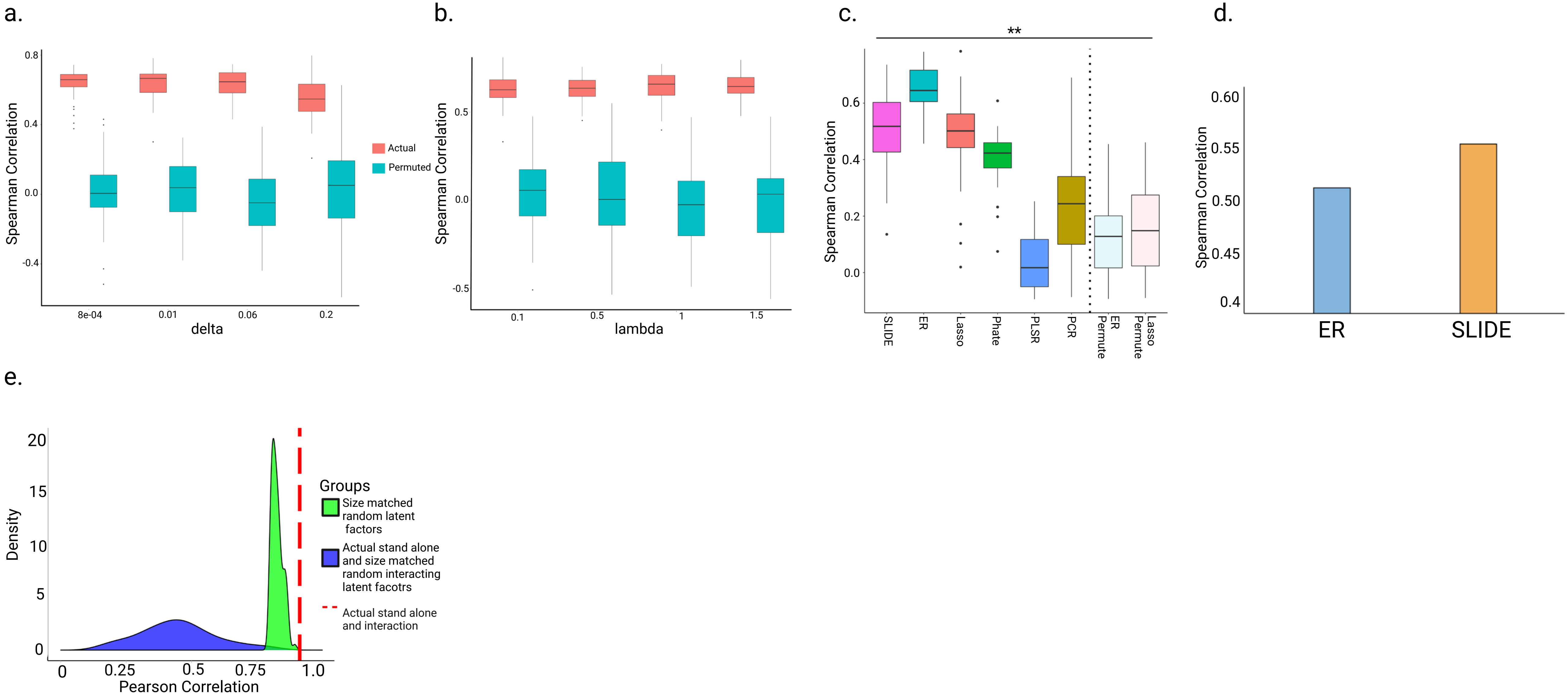

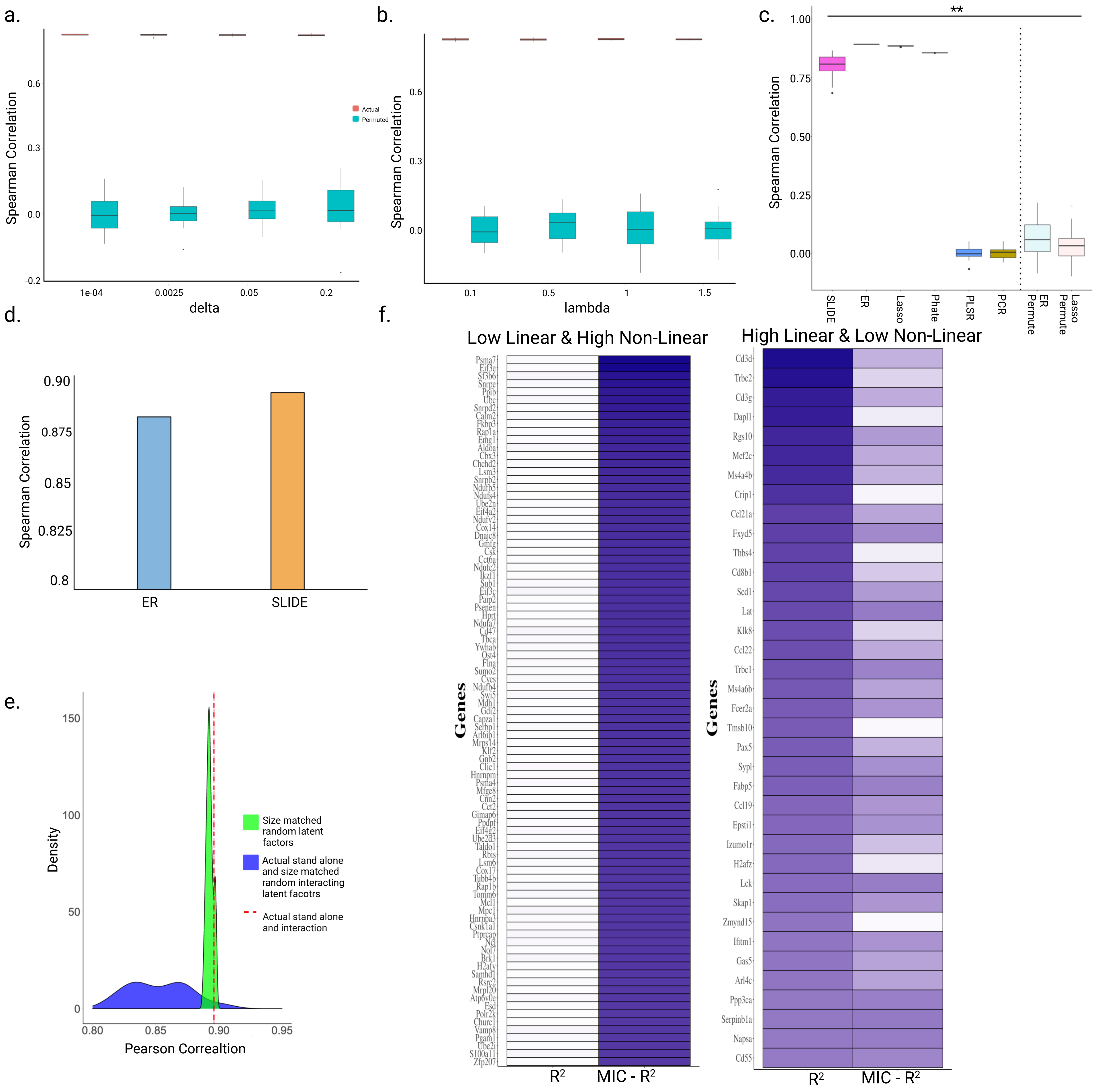

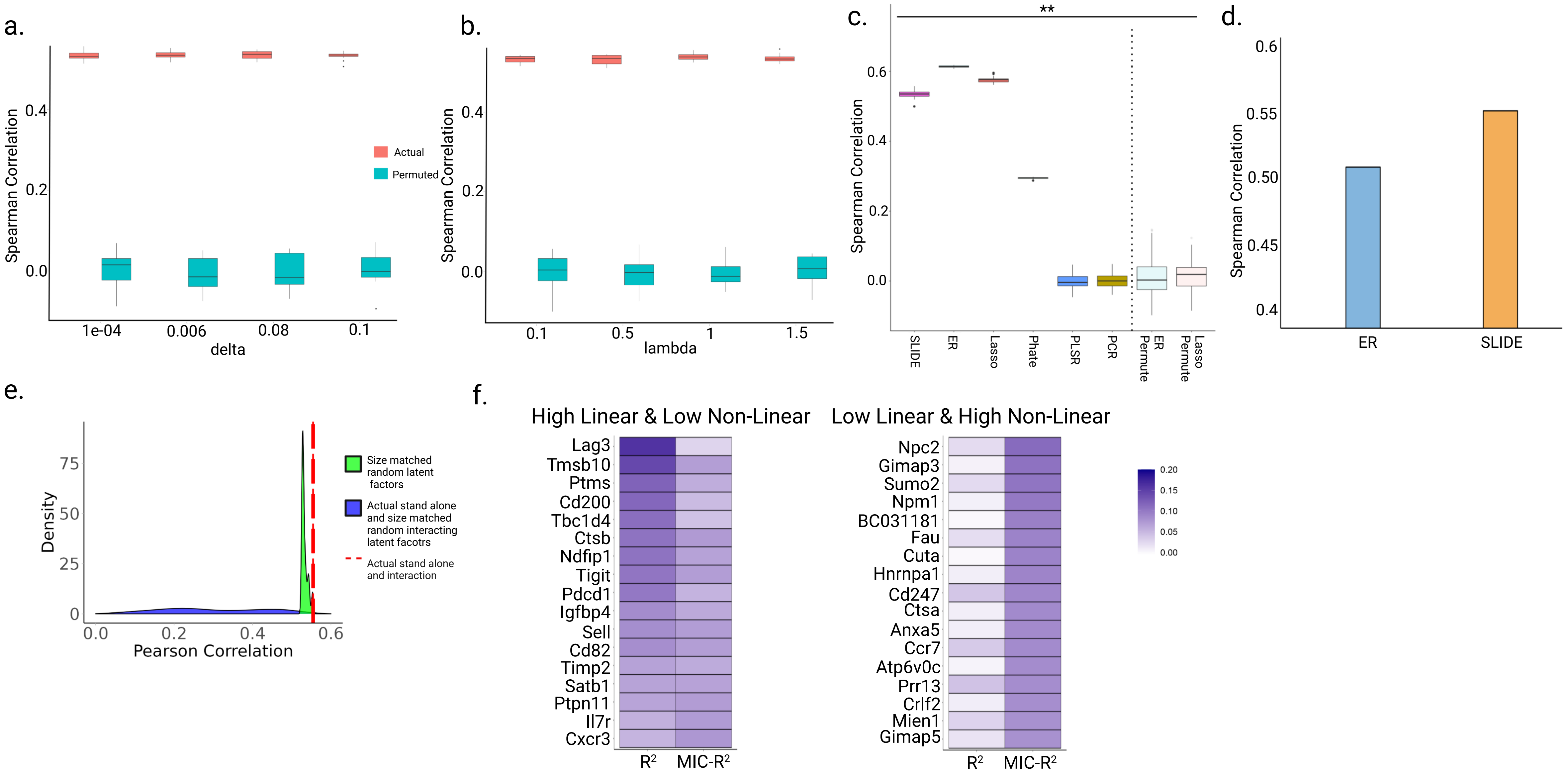
