## Supplementary Methods for "SLIDE: Significant Latent Factor Interaction Discovery and Exploration across biological domains"

Table 1: Notation.

| Notation | Description |
|---|---|
| $SFList$ | list of the selected feature |
| $SFreq$ | Selected Feature Frequency |
| $KnF$ | Filter Frequency Selected Features by knockoff |
| $Nint$ | Number of iterations |
| $L$ | Number of partition |
| $F$ | Feature Split Size |
| $K$ | Number of latent variables |
| $Z$ | The latent variables |
| $y$ | The dependent variable |
| $n$ | Number of samples |
| $PS$ | Potential Stand Alone Variables |
| $FSA$ | Final Stand Alone |
| $IntF_i$ | Interaction Feature i |
| $SIntF_i$ | Selected Interaction for Stand Alone Feature $i$ |
| $q_l$ | l-th Partition of Z matrix |
| $spec$ | The frequency cutoff |

**Algorithm 1:** Repeatedknockoff

**input** : $Z$ of size $n \times K$, spec

**output:** $KnF$

**for** $i \leftarrow 1$ *to* $Nint$ **do**

  | $SFList[i] \leftarrow Knockoff(Z, y)$

**end for**

$SFreq = ConstructFrequency(SFList)$

$KnF = Filter(SFreq, spec)$

Because the data is high-dimensional, we partitioned the features into a smaller set of features with $L = \lceil \frac{p}{F} \rceil$ number of sets.

we defined the partitioned Z data as $q_l$ using following equation:

$q_l = Z[, (l-1)F + 1, (l-1)F + 2, \ldots, lF]$ where $l \in \{1, 2, .., L\}$

we regressed the latent variable $Z_i$ and its interactions on $y$, and used the predict value as the embedding $\hat{E}_i = Z_i\hat{\beta}_i + \Sigma_{i,j\in FSA}\hat{\beta}_{ij} Z_i Z_{i|j}$ ($i|j$:j-th interaction selected for i-th variable) interactors by SLIDE alogithm. The $\hat{\beta}$ are calculated as least square estimate: $\hat{\beta}_{\mathbf{i}} = (Z^T_{i|j,i\in FSA,j\in SIntF_i} Z_{i|j,i\in FSA,j\in SIntF_i})^{-1} Z^T_{i|j,i\in FSA,j\in SIntF_i} y$

**Algorithm 2:** SLIDE

**input :** $Z$ matrix of size $n \times K$, spec, $y$
**output:** $Zi$ and $SIntZi$ for i $\in FSA$

**Stage 1 : Finding potential Standalone variable**

**for** $l \leftarrow 1$ **to** $L$ **do**
  $q_l = Z[, (l-1)F + 1, (l-1)F + 2, \ldots, lF]$
  $PS[l] \leftarrow Repeatedknockoff(q_l, y, spec)$
**end for**

**Stage 2: Finding Significant standalone latent factor**

$FSA < -Repeatedknockoff(PS, y, spec)$

**Stage 3: Finding significant interacting latent factor of each significant standalone**

**for** $i \in FSA$ **do**
  $\mathrm{IntZ}_i \leftarrow GetInteraction(\mathrm{Z}_i,\ Z)$
  $yc_i \leftarrow y - Z_i({Z_i}^T Z_i)^{-1} {Z_i}^T y$
  $SIntZ_i \leftarrow Repeatedknockoff(IntZ_i, yc_i, spec)$
**end for**
